# Higher phage virulence accelerates the evolution of host resistance

**DOI:** 10.1101/2021.03.26.437141

**Authors:** Carolin C. Wendling, Janina Lange, Heiko Liesegang, Michael Sieber, Anja Pöhlein, Boyke Bunk, Jelena Rajkov, Henry Goehlich, Olivia Roth, Michael A. Brockhurst

**Affiliations:** GEOMAR Helmholtz Centre for Ocean Research Kiel, Marine Evolutionary Ecology, Düsternbrooker Weg 20, 24105 Kiel, Germany; ETH Zürich, Institute of Integrative Biology, Universitätstrasse 16, CHN D 33, 8092 Zürich, Switzerland; Georg-August-University Göttingen, department of genomic and applied microbiology, Grisebachstr 8 37077 Göttingen, Germany; Max Planck Institute for Evolutionary Biology, August-Thienemann-Str. 2, 24306 Plön, Germany; Leibniz Institute DSMZ-German Collection of Microorganisms and Cell Cultures, Department Bioinformatics and Databases, Inhoffenstr. 7B, 38114 Braunschweig, Germany; Kiel University, Marine Evolutionary Biology, Am Botanischen Garten 1-9, 24118 Kiel, Germany; Division of Evolution and Genomic Sciences, University of Manchester, Dover Street, Manchester M13 9PT, UK

**Keywords:** virulence, filamentous phages, experimental evolution, resistance evolution

## Abstract

Parasites and pathogens vary strikingly in their virulence and the resulting selection they impose on their hosts. While the evolution of different virulence levels is well studied, the evolution of host resistance in response to different virulence levels is less understood and as of now mainly based on observations and theoretical predictions with few experimental tests. Increased virulence can increase selection for host resistance evolution if resistance costs are outweighed by the benefits of avoiding infection. To test this, we experimentally evolved the bacterium *Vibrio alginolyticus* in the presence of two variants of the filamentous phage, VALGΦ8, that differ in their virulence. The bacterial host exhibited two alternative defence strategies against future phage infection: (1) super infection exclusion (SIE) whereby phage-infected cells were immune to subsequent infection at a cost of reduced growth, and (2) surface receptor mutations (SRM) in genes encoding the MSHA type-IV pilus providing resistance to infection by preventing phage attachment. While SIE emerged rapidly against both phages, SRM evolved faster against the high virulence compared to the low virulence phage. Using a mathematical model of our system we show that increasing virulence strengthens selection for SRM due to the higher costs of infection suffered by SIE immune hosts. In both the experiments and the model, higher levels of SRM in the host population drove more rapid phage extinction. Thus, by accelerating the evolution of host resistance, more virulent phages caused shorter epidemics.

## INTRODUCTION

Infectious organisms vary strikingly in their level of virulence and the resulting selection they impose on hosts. Indeed, even closely related viruses, such as different strains of myxoma (Fenner and Marshall 1957) or corona viruses (Weiss and Leibowitz 2011), can differ greatly in virulence. While the evolution of virulence has been studied extensively during the last two decades, both using selection experiments (Bull, Molineux et al. 1991, Turner, Cooper et al. 1998, Messenger, Molineux et al. 1999) and observations of parasites evolved in nature (Herre 1993, Ebert 1994), how hosts respond to virulence-mediated selection is less well-explored. Our understanding of how virulence will impact evolutionary trajectories of resistance in a host population, and how these trajectories change with different levels of virulence, is mainly based on observational patterns (Kraaijeveld and Godfray 1999, Gates, Staley et al. 2021) and theory (van Baalen 1998, Boots and Haraguchi 1999, Restif and Koella 2003) with few experimental tests (Kraaijeveld and Godfray 1997). In general, increased virulence strengthens selection for the evolution of host resistance if the costs of resistance are outweighed by the benefits of avoiding infection (van Baalen 1998, Boots and Haraguchi 1999, Restif and Koella 2003). As such, at very low virulence, although infection is common, resistance is not favoured because the cost of resistance is likely to exceed any benefits of avoiding mild disease (Restif and Koella 2003). With increasing virulence, resistance is more strongly selected as the cost of resistance becomes outweighed by the detrimental effects of more severe disease, leading to the more rapid evolution of resistance (van Baalen 1998). However, at extreme levels of high virulence, selection for resistance can weaken once more, due to declining infection prevalence (Boots and Haraguchi 1999). Experimental tests of these predictions are, however, rare.

To explore how different levels of virulence influences the dynamics of host resistance evolution, we designed a selection experiment using the model bacterium *Vibrio alginolyticus* K01M1 as a host and two variants of the filamentous phage, VALGΦ8, that differ in their virulence but are otherwise isogenic ((Chibani, Hertel et al. 2020), Table 1). Filamentous phages (family *Inoviridae*)—i.e., long, thin proteinaceous filaments which contain a circular single-stranded DNA genome—have been shown to be ideal model systems to study virulence evolution (Bull, Molineux et al. 1991, Messenger, Molineux et al. 1999). Filamentous phages establish chronic infections whereby virions are continuously released without lysis. Although filamentous phages do not kill their host, infections often result in reduced host growth rates (Mai-Prochnow, Hui et al. 2015, Hay and Lithgow 2019). This reduction in growth can result from an overexpression of gI, which can even result in cell death or from phage-encoded proteins inserted into the bacterial membrane (Mai-Prochnow, Hui et al. 2015). Thus, we define virulence here as the reduction in bacterial growth resulting from phage infection, which we can directly quantify by measuring the reduction in bacterial growth rate caused by phage infection relative to the growth rate of phage-free cultures.

**Table 1.**
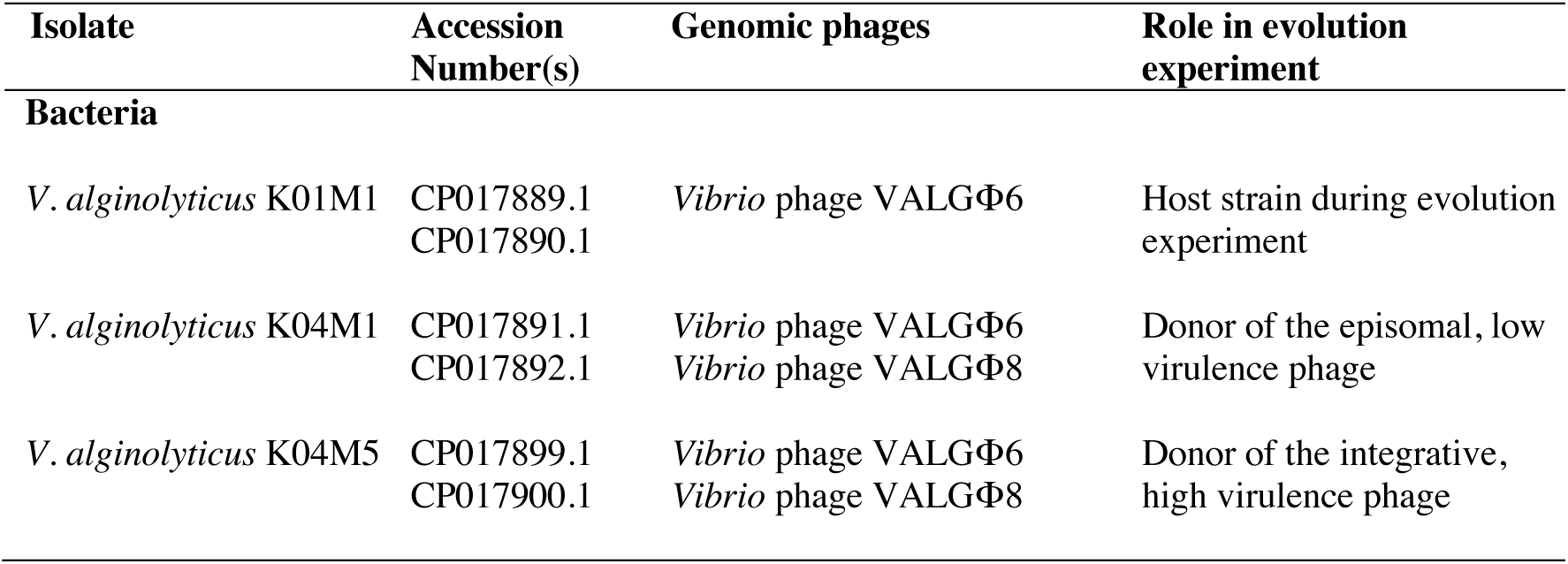
Strains (including NCBI accession numbers for chromosome 1 and 2) used in the present study

| Isolate | Accession Number(s) | Genomic phages | Role in evolution experiment |
| --- | --- | --- | --- |
| **Bacteria** |  |  |  |
| *V. alginolyticus* K01M1 | CP017889.1<br>CP017890.1 | *Vibrio* phage VALGΦ6 | Host strain during evolution experiment |
| *V. alginolyticus* K04M1 | CP017891.1<br>CP017892.1 | *Vibrio* phage VALGΦ6<br>*Vibrio* phage VALGΦ8 | Donor of the episomal, low virulence phage |
| *V. alginolyticus* K04M5 | CP017899.1<br>CP017900.1 | *Vibrio* phage VALGΦ6<br>*Vibrio* phage VALGΦ8 | Donor of the integrative, high virulence phage |

**Table 2.** Parameter values of mathematical model and their biological meaning

| Parameter | Biological meaning | Value |
| --- | --- | --- |
| $r$ | Maximum growth rate (mgr) of ancestor $B$ | 2.5 ($\text{h}^{-1}$) |
| $K$ | Carrying capacity of bacteria | $10^9$ cells/ml |
| $w$ | Curvature parameter | 0.02 |
| $v$ | Virulence | variable |
| $\phi$ | Phage adsorption rate | $10^{-8}$ ($\text{h}^{-1}$) |
| $\beta$ | Phage production rate | 50 (phages/cell $\text{h}^{-1}$) |

During chronic infections, most phage genes are repressed to ensure host cell viability (Bondy-Denomy and Davidson 2014). This is achieved through the action of prophage encoded repressor proteins which also prevent superinfection i.e., superinfection exclusion (SIE) by the same or closely related (Refardt 2011) phage(s). Many filamentous phages, including vibriophages from the present study, can provide SIE immunity through the production of the phage-encoded receptor-binding protein pIII which blocks primary and secondary phage receptors (Mai-Prochnow, Hui et al. 2015). As such, chronically infected host cells become protected from subsequent infection through SIE. Alternatively, it is possible for bacteria to acquire resistance to filamentous phage infection through mutations causing alterations to the surface receptors that the phages bind to, thus preventing phage infection (Jouravleva, McDonald et al. 1998). How phage virulence alters selection for SIE versus surface receptor modification (SRM) resistance is unclear.

Combining experimental evolution with whole genome sequencing, we show that SIE immunity arose rapidly and at a similar rate against both phages, whereas SRM evolved more rapidly against the high compared to the low virulence phage, driving faster extinction of the high virulence phage. Using an experimentally parameterised mathematical model we show that accelerated replacement of SIE immunity by SRM was driven by increasing costs of infection, in terms of reduced growth, suffered by SIE immune hosts with increasing phage virulence. Resistance mutations were identified in genes encoding the MSHA type IV pilus, which pleiotropically caused reduced motility of these resistant bacteria. Together these data show that higher phage virulence accelerated the evolution of resistance, which consequently drove faster phage extinctions and shorter epidemics.

## RESULTS

### Ecological dynamics vary according to phage virulence

To explore how variation in virulence influences the dynamics of host resistance evolution, we experimentally evolved the bacterium *Vibrio alginolyticus* K01M1 with or without one of two isogenic filamentous phages that differ in their virulence—VALGΦ8_K04M5_ which reduces bacterial growth by 73% (higher virulence) or VALGΦ8_K04M1_ which reduces bacterial growth by 58% (lower virulence, Table 1)—for 30 serial transfers (∼240 bacterial generations). We first compared the ecological dynamics of bacterial and phage populations between treatments. Phages reduced bacterial densities by several orders of magnitude in both phage treatments compared to no phage control populations (Figure 1a). The immediate reduction (measured 24 hours post infection [hpi]) in bacterial density was greater in populations exposed to the higher virulence phage (VALGΦ8_K04M5_) than the lower virulence phage (VALGΦ8_K04M1_; Figure 1a). Correspondingly, in both treatments, phages amplified massively and rapidly, reaching 3.01×10^12^ PFU/ml (VALGΦ8_K04M5_) 24 hpi and 2.83×10^12^ PFU/ml (VALGΦ8_K04M1_) 48 hpi (Figure 1b), before declining to levels comparable to control populations (note that the genome of *V. alginolyticus* K01M1 contains a resident phage, VALGΦ6, that produces phage particles at a low background rate). These data suggest that the strong reduction in bacterial densities at the beginning of the experiment (Figure 1a) directly resulted from the costly production of viral particles (Figure 1b). Over time, however, the densities of bacterial populations exposed to the higher virulence phage recovered three times faster than populations exposed to the lower virulence phage (significant phage:transfer interaction in gls-model: F_15,186_=6.58, p<0.001, Figure 1a). Bacterial population recovery was accompanied by declining phage densities in both treatments, but phage survival varied according to phage virulence (log-rank test: Chisq_1_=4.9, p=0.03), with the higher virulence phage going extinct more rapidly than the lower virulence phage (Figure 4a).

**Figure 1.**
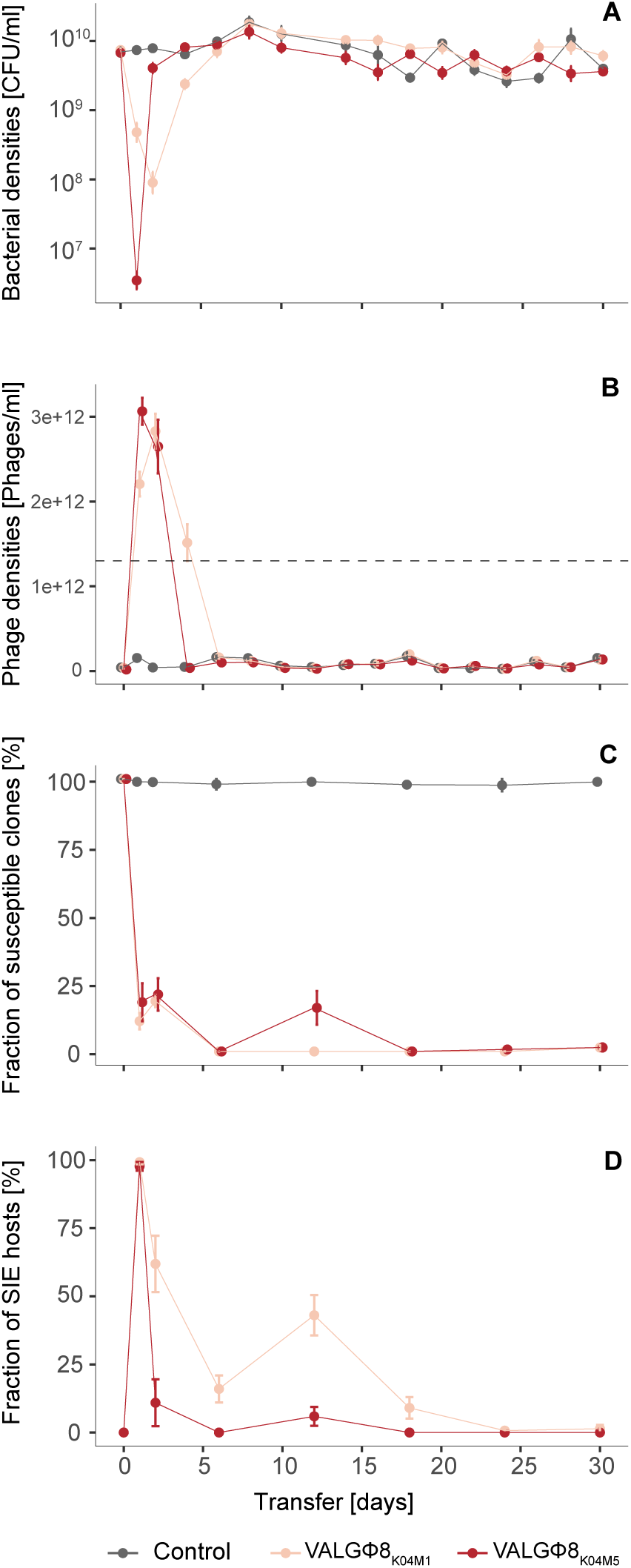
Population dynamics over 30 transfers. (A) Bacteria in CFU/ml, (B) Phages in PFU/ml, the grey dashed line represents the quantification limit below which quantifying filamentous phages using spectrophotometry is inaccurate, note: free phages in the control treatment stem from the low-replicating resident phage VALGΦ6 (see Table 1) (c) Fraction of susceptible clones (n=24), and (d) Fraction of SIE hosts within phage-resistant clones. Fractions are based on 24 random clones per replicate population per timepoint. In all panels, data are represented as mean of six replicate populations per treatments, error bars represent standard errors. Colours correspond to one of three experimental treatments, lower virulence VALGΦ8_K04M1_ (light red), higher virulence VALGΦ8_K04M5_ (dark red), no phage (grey).

### Rapid emergence of superinfection exclusion immunity

These bacteria-phage population dynamics suggest that the emergence of bacterial defences against phage infection may have enabled recovery of the host population. Consistent with this hypothesis, the proportion of susceptible hosts rapidly declined to zero within 24 hours in both treatments and remained so for the duration of the experiment (Figure 1c). Bacteria can develop protection from filamentous phage infection by two distinct mechanisms: superinfection exclusion (SIE) immunity, where already infected cells are protected from subsequent infection by the same phage through phage-encoded genes (Susskind, Wright et al. 1974, Uc-Mass, Loeza et al. 2004, Sun, Gohler et al. 2006, Cumby, Edwards et al. 2012), or resistance, for instance via modification of the bacterial phage receptor, preventing phage from entering the host cell (Jouravleva, McDonald et al. 1998). To quantify the frequency of SIE immunity we used PCR with primers that target specifically VALGΦ8 to test for the presence of the relevant phage in the bacterial genome (the presence of a PCR product suggests SIE due to the presence of VALGΦ8). SIE rapidly increased in frequency and dominated bacterial populations in both treatments after 24 hours (Figure 1d). However, after 48 hours, the proportion of SIE hosts began to decline, and did so significantly faster in populations that had been exposed to the higher virulence phage (Figure 1d, significant phage:transfer interaction in glm: F_6,60_=10.18, p<0.001). Given that these populations contained no susceptible bacteria from 24 hours onwards (out of 24 tested colonies per timepoint), the subsequent decline of SIE hosts suggests their displacement by the invasion of resistant genotypes, and that this was more strongly selected for by the higher virulence phage.

### Resistance is associated with mutations in MSHA type IV pilus encoding genes

To test if the decline of SIE hosts after 24 hours was driven by the invasion of surface receptor modification (SRM) resistance, we used whole genome sequencing (WGS) of two randomly chosen clones from each population isolated at transfer 2: one PCR-positive clone (SIE) and one PCR-negative clone (resistant but not phage carrying) to identify mutations. We observed no loci with mutations on chromosome 2 or the plasmid pl9064. However, on chromosome 1 we identified 12 loci with mutations that were not present in clones from the control treatment, suggesting that these were associated with phage-mediated selection. Of these 12 loci, two were affected in PCR-positive and PCR-negative clones. This included an intergenic region between tRNA-Val and the 23S ribosomal RNA, that has been repeatedly hit in both clone-types and phage-treatments, but whose function we cannot explain. The remaining ten loci were exclusive to PCR-negative clones suggesting a potential role in evolved phage resistance. Of these nine loci, eight had substitutions, duplications, insertions, or deletions in four different genes belonging to the MSHA type IV pilus operon (*mshL, mshE, mshG, K01M1_28150;* Figure 2a/ Table S1). Among those, three caused severe frameshift mutations that presumably have a high impact on the function of these proteins. While the locus (*K01M1_28150*) was affected twice in both phage treatments, mutations in *mshL* and *mshE* occurred exclusively in response to the higher virulence phage and mutations in *mshG* in response to the lower virulence phage. Moreover, we found more mutated MSHA-loci among clones exposed to the higher virulence (5/6) compared to the lower virulence phage (3/6). This supports of our previous findings, which suggested a stronger selection for resistance against the higher virulence phage.

**Figure 2.**
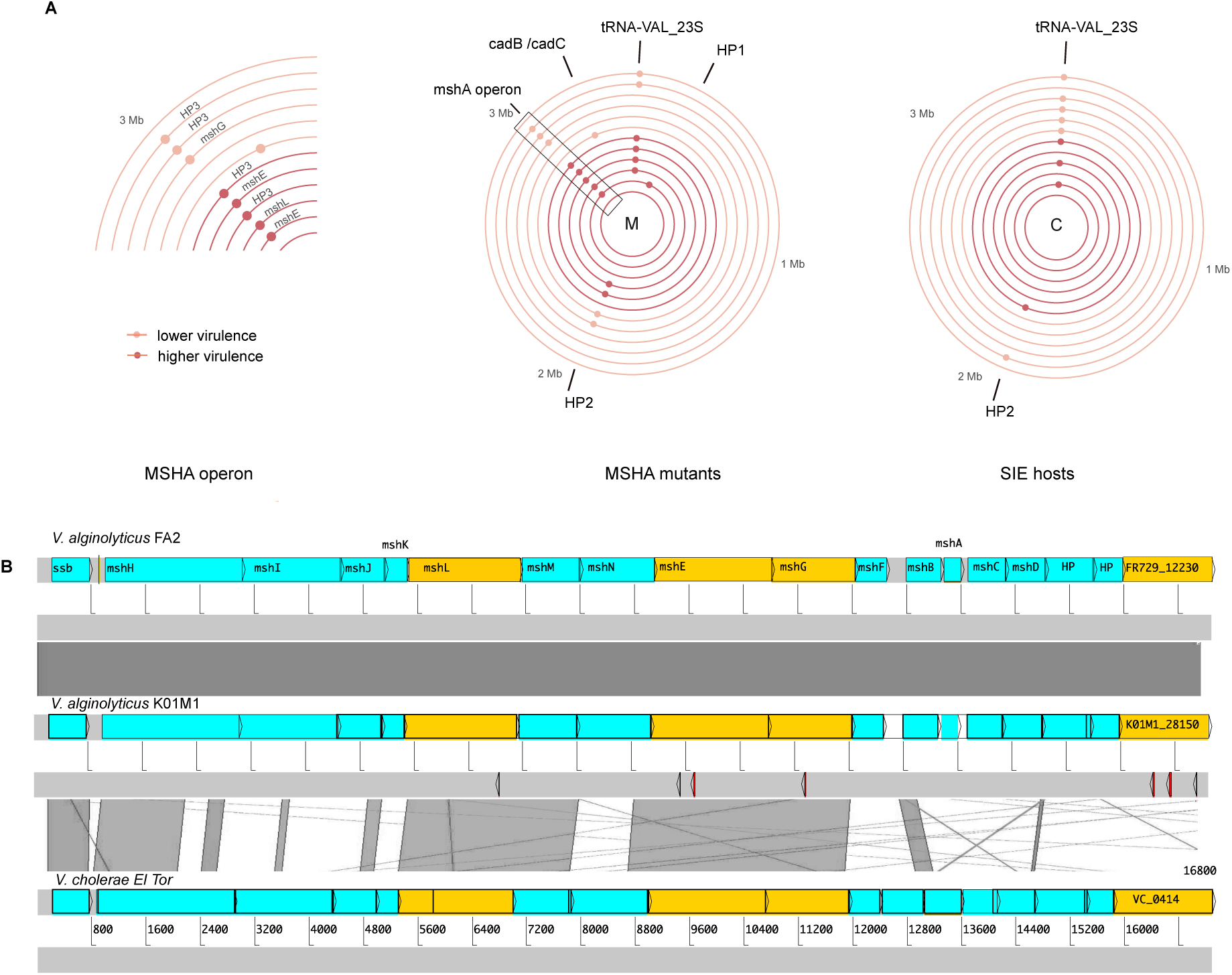
(A) Genetic loci on chromosome 1 under positive selection as indicated by parallel genomic evolution in populations exposed to phages: right: SIE hosts; middle: SRM hosts; left: zoom into MSHA-operon region from SRM hosts. Only loci which are not present in control populations are shown. Concentric circles correspond to one clone isolated from either the higher virulence VALGΦ8_K04M5_ (six inner circles, dark red) or the lower virulence VALGΦ8_K04M1_ phage (six outer circles, light red). Each coloured point corresponds to one mutation event on the respective clone. HP = hypothetical protein; HP3 corresponds to locus tag K01M1_28150. For more detailed information on the underlying mutation see Table S1. (B) Structure of the MSHA-operon and comparative genomics comprising MSHA-operons from *V. alginolyticus* FA2 (top), *V. alginolyticus* K01M1, and *V. cholerae* El Tor (bottom). Similarity between regions is indicated by dark grey blocks, genes with detected mutations are marked in orange, detected mutations are marked as arrows below *V. alginolyticus* K01M1.

The absence of mutated MSHA-loci in PCR-positive clones paired with a high prevalence in PCR-negative clones (8/12) suggests strongly parallel evolution of phage resistance. The MSHA operon is highly conserved across *Vibrio* clades (Figure 2b), and we found one corresponding ortholog to each gene in the *V. cholerae* El Tor MSHA operon (Figure 2b). This suggests that, similar to other vibrios (Jouravleva, McDonald et al. 1998), the MSHA type IV pilus plays an important role in resistance against VALGΦ8. Note, a search of all assembled genomes for CRISPR associated genes as well as for CRISPR array like repetitive sequence patterns did not yield any results. All PCR-negative phage resistant clones are from here onwards referred to as surface receptor mutant SRM hosts. The genomic data also confirmed that clones with a positive PCR result (i.e., SIE host) all contained the respective phage genome, which did not integrate into the chromosome but existed episomally in all sequenced clones (Table S2; Figures S3).

We found four PCR negative clones that were resistant to infections with ancestral phages but did not acquire mutations within the MSHA operon. One explanation could be phenotypic resistance, where phage adsorption to bacteria is strongly reduced (Bull, Vegge et al. 2014). Another explanation could be inactivation of genes required for phage replication (Martinez and Campos-Gomez 2016). For instance, we found two PCR-negative clones with mutations in hypothetical proteins whose functions we do not know.

### Virulence determines the rate of resistance evolution in a mathematical model

To generalize our findings across a wider range of virulence levels we developed an experimentally parameterized mathematical model. As in the experiment, bacterial densities dropped by several orders of magnitude upon phage infection (Figure 3a). By simulating the infection dynamics over a wider range over virulence levels, we found that this drop occurred later and was less strong with decreasing virulence. While phage densities, irrespective of virulence, peaked 24 hpi, phages persisted longer and at higher levels when they were less virulent (Figure 3b). Similar to the experiment, the model predicts that SIE immunity emerges rapidly within 24 hpi (Figure 3c) but will only reach high levels if virulence is < 1. To capture the displacement of SIE by SRM hosts we implemented a cost of reduced growth for SIE hosts which is directly linked to virulence (Figure 4c), i.e., the higher the virulence of the infecting phage, the lower the growth rate of the SIE host. SRM hosts grew at the same rate as the non-resistant clones (Figure 4e).

**Figure 3.**
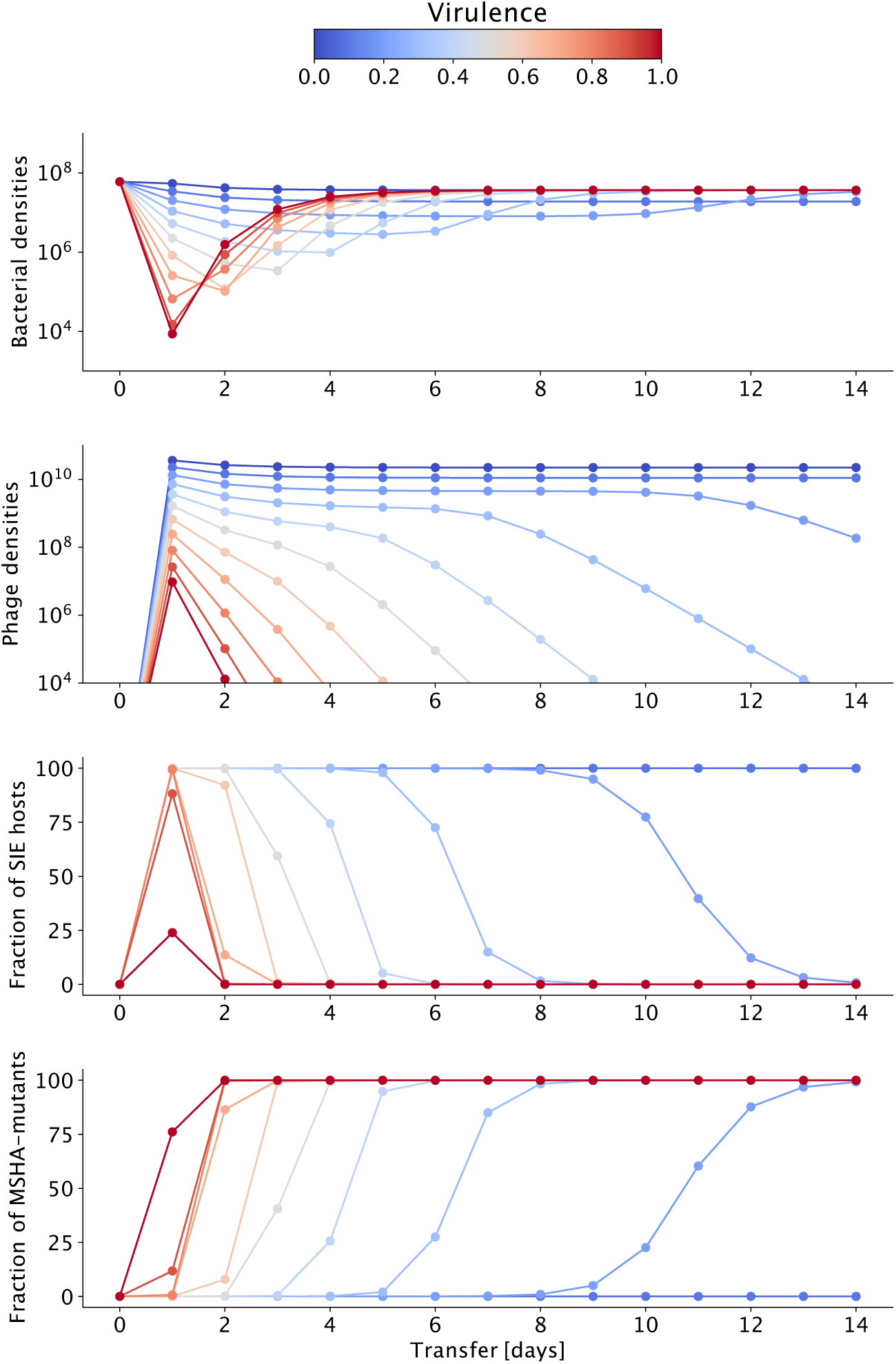
Results of model simulations of 14 transfers for (A) Bacteria in CFU/ml, (B) Phages in PFU/ml, (C) SIE hosts, and (D) SRM hosts depending on phage virulence (colour coded from blue: no virulence to red: high virulence).

**Figure 4.**
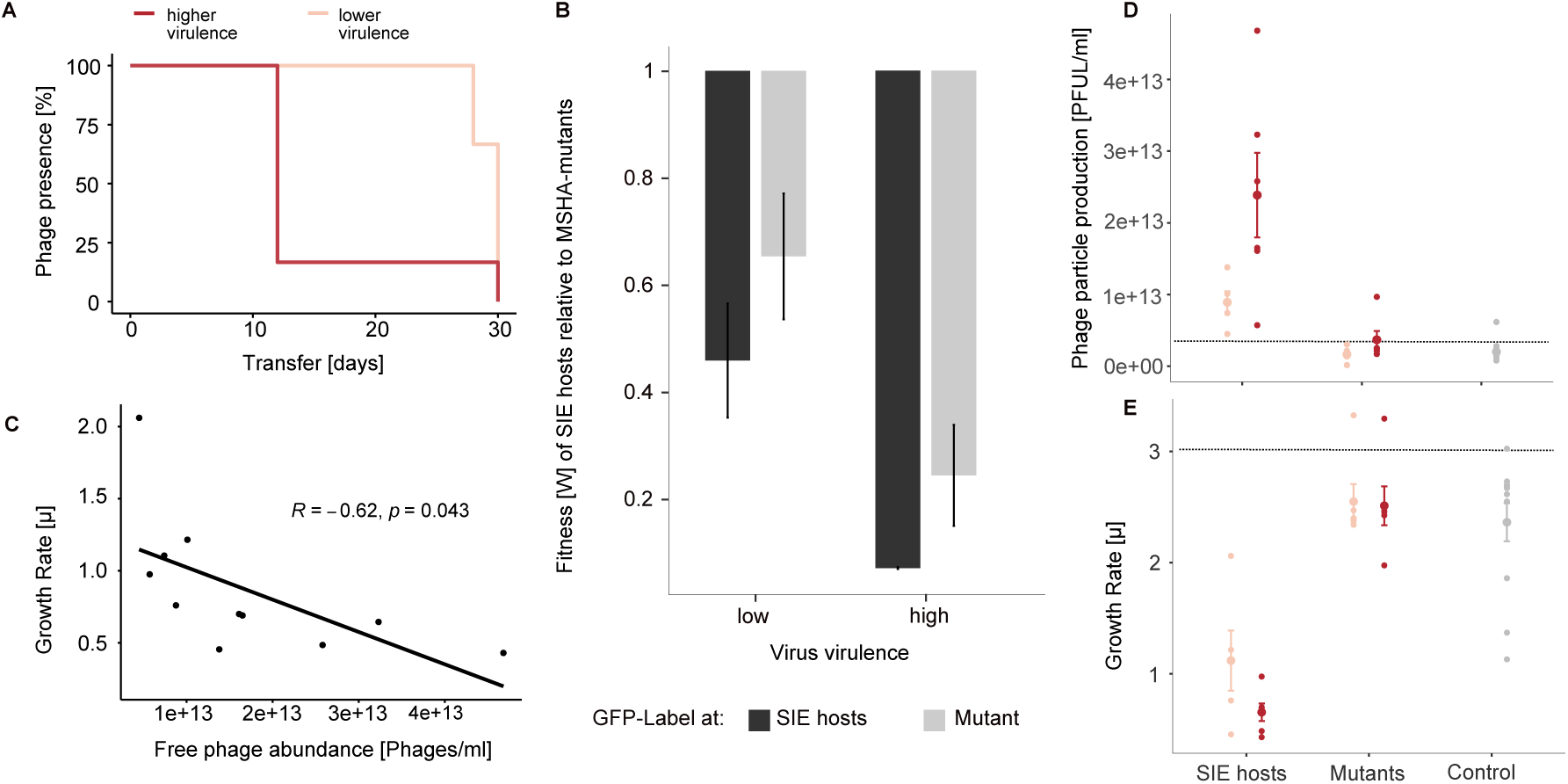
Phage prevalence (a) and fitness effects of evolved phage resistance versus immunity (b-e): (a) Phage prevalence for each co-evolving population in the presence of higher virulence phage VALGΦ8_K04M5_ (dark red) or the lower virulence phage VALGΦ8_K04M1_ (light red) over 30 transfers. (b) Darwinian fitness of SIE relative to SRM hosts. A value of one corresponds to equal fitness. To account for potential costs associated with the GFP protein, competitions were performed where either the SIE or the SRM host were labelled (n=3). (c) Correlation between bacterial growth rate [*µ*] and production of free phages measured as PFU/ml per clone. (d) Phage particle production [PFU/ml] and (e) Growth rate *µ*: both measured after 24 hours of bacterial growth for SIE hosts, SRM hosts, clones from the control populations (grey), and the ancestral K01M1 strain (horizontal line). Clones exposed to lower virulent VALGΦ8_K04M1_ are shown in light red, clones exposed to higher virulent VALGΦ8_K04M5_ in dark red. Phages from the ancestral K01M1, from SRM hosts and the control clones stem from an ancestral filamentous *Vibrio* phage VALGΦ6 integrated on chromosome 2 of K01M1 (Table 1). Shown are means ± SEM, n=24.

We found that SRM hosts increased and replaced SIE hosts faster with increasing levels of virulence (Figure 3 c,d). Our model shows that this replacement occurs across a wide range of virulence levels, which we were not able to capture in the experiment. The faster replacement of SIE by SRM hosts at higher virulence levels is driven by higher costs i.e., reduced growth of infection in SIE hosts, which increase monotonically with increasing phage virulence. Overall, our simulations predict that selection for resistance increases with virulence and that this is directly related to the costs of SIE, and thus resistance is more likely to evolve against higher virulence infections.

### Relative fitness of surface receptor mutants increases with phage virulence

To directly test our model predictions, that the fitness benefit of SRM relative to SIE immunity increases with increasing phage virulence, we performed a pairwise competition experiment, in which we quantified the relative fitness of SRM against SIE hosts (Figure 4b). The fitness benefit of the resistance mutation was higher against bacteria carrying the higher virulence phage compared to bacteria carrying the lower virulence phage (significant treatment term in linear model with treatment, GFP-label and the interaction thereof as fixed factors: F_1,8_=18.63, p=0.003, Table S3). These fitness data, which are consistent with the more rapid decline of VALGΦ8_K04M5_-carriers than VALGΦ8_K04M1_-carriers observed in the selection experiment and consistent with model predictions suggest stronger selection for SRMs when exposed to a higher virulence phage. This explains the dynamics of the SIE hosts in the selection experiment, which went to extinction in five out of six populations exposed to the higher virulence phage 12 days post infection but were able to persist until the end of the experiment, i.e., transfer 30, albeit at very low frequencies, in five out of six populations exposed to the lower virulence phage.

Bacterial population densities during the selection experiment were negatively correlated with the number of SIE hosts per population (Pearson’s correlation without zero inflation Φ-K04M1: r=0.69, t_21_=-4.38, p<0.001, Φ-K04M5: r=0.92, t_7_=-6.29, p<0.001; Figure S4). This implies that, even though the majority of the clones in the populations were protected from further infection, bacterial populations were unable to recover as long as the dominating mechanism of defence was SIE immunity, presumably due to virulence, resulting from the strong reduction in bacterial growth rate. To test this, we quantified differences in phage production and tested if phage production impaired bacterial growth in SIE hosts. VALGΦ8_K04M5_-carriers produced approximately 3.5 times more phage particles than VALGΦ8_K04M1_-carriers (VALGΦ8_K04M5_: mean = 2.39×10^13^ PFU/ml ± 1.44×10^13^, VALGΦ8_K04M1_: mean = 8.92×10^12^ PFU/ml ± 3.43×10^12^, Figure 4d), and phage production significantly impaired bacterial growth (significant negative correlation between the amount of produced phages and bacterial growth rate, Figure 4c). Direct comparisons of growth rates among resistant clones showed that SIE hosts also grew slower than SRM hosts (VALGΦ8_K04M5_: paired *t*-test: t_6.93_=-9.69, p<0.001; VALGΦ8_K04M1_: paired *t*-test: t_6.5_=-4.58, p=0.003, Figure 4e). Together, these data suggest that SIE buys time, offering protection against subsequent infection, but at the cost of suffering the virulence of being infected. As in the model, where we predicted that the costs of SIE increase monotonically with phage virulence, SIE is eventually replaced by SRM, which happens faster with increasing levels of virulence, where the fitness benefits of SRM are greater. Ultimately dominance of host populations by SRM hosts resulted in faster extinction of higher virulence phages, which were unable to overcome evolved host resistance.

### Resistance leads to secondary costs

Lastly, we tested whether the observed mutations in the MSHA-pilus genes impair bacterial motility. We observed reduced swimming motility of SRM hosts compared to ancestral bacterial strains (Video Supplementary material).

## DISCUSSION

Theory predicts that increasing virulence results in stronger selection for host resistance if the benefits of avoiding infection are higher than the costs of resistance (van Baalen 1998, Boots and Haraguchi 1999, Restif and Koella 2003), but experimental data are rare (Kraaijeveld and Godfray 1997). Using two filamentous phages that differ in their virulence in a selection experiment, we found that SIE immunity arose rapidly and at a similar rate irrespective of phage virulence. In contrast, SRM, which replaced SIE immunity, evolved more rapidly against the high compared to the low virulence phage. Our mathematical modelling confirmed that increasing virulence strengthens selection for SRM because SIE immunity becomes more costly with increasing virulence. Thus, higher levels of SRMs in the host population caused faster phage extinction and ultimately shorter epidemics.

Resistance arose from mutations in genes encoding the MSHA type IV pilus, a common receptor for filamentous phages (Hay and Lithgow 2019). These mutations reduced the motility of resistant clones suggesting pleiotropic fitness costs of phage resistance mutations. Such secondary costs, i.e., reduced motility or even pathogenicity are commonly observed during phage resistance evolution involving multifunctional structures on bacterial cell surfaces, for a review, see (Leon and Bastias 2015). In many natural environments, motility is an important pathogenicity factor (Proft and Baker 2009), which is often crucial for the establishment of acute infections but less important during chronic infections of eukaryotic hosts. A prominent example are infections of cystic fibrosis (CF) lungs by *Pseudomonas aeruginosa*. While initial colonizers are motile, adaptation to CF lungs during chronic infections is often characterized by loss of motility (Wong, Rodrigue et al. 2012, McElroy, Hui et al. 2014). Thus, we predict that the replacement of SIE hosts by non-motile SRM hosts may be constrained to laboratory environments but may also occur in biofilms, where such antagonistic pleiotropic costs of surface receptor modifications are lower than the costs of SIE. In contrast, selective pressures occurring in eukaryotic hosts, such as resource competition, might reverse this effect and explain why filamentous phages, including highly virulent versions of VALGΦ8 persist in environmental isolates (Chibani, Hertel et al. 2020).

Filamentous phages are very common features of bacterial genomes (Roux, Krupovic et al. 2019), including those of environmental *Vibrio* strains closely related to our model strain K01M1, of which all carry VALGΦ6 and more than 50% VALGΦ8 (Chibani, Hertel et al. 2020). While incorporating filamentous phage genomes into their own genome (but not necessarily into the bacterial chromosome) provides bacteria with immunity to future infection—through SIE immunity mediated by phage-encoded genes—we show that this comes at a fitness cost that scales positively with the virulence of the phage. Higher phage virulence selects for faster replacement of SIE immunity with SRM, causing phage extinction (Figure 4a). Thus, our data suggest, that to be able to establish long-term chronic infections, filamentous phages must either evolve very low levels of virulence (Lerner and Model 1981), such that the resulting cost of virulence is outweighed by the cost of resistance mutations, or alternatively they must contribute positively to bacterial fitness by providing beneficial ecological functions (Hay and Lithgow 2019). Those benefits may derive either from phage-encoded gene functions e.g., toxins (Waldor and Mekalanos 1996, Gonzalez, Lichtensteiger et al. 2002, Rice, Tan et al. 2009), or from properties of the phage particles themselves e.g., forming the biofilm extracellular-matrix (Secor, Sweere et al. 2015), or acting as decoys for the vertebrate immune response (Sweere, Van Belleghem et al. 2019). Phage-mediated fitness benefits are often environmentally dependent (Gonzalez, Lichtensteiger et al. 2002, Derbise, Chenal-Francisque et al. 2007, Chouikha, Charrier et al. 2010, Wendling, Refardt et al. 2020) and the prevalence of filamentous phages in bacterial genomes is higher in those isolated from eukaryotic infections, where filamentous phages often encode important enterotoxins, than those isolated from natural environments (Mai-Prochnow, Hui et al. 2015). Even though we have not yet identified any associated ecological benefits, the high prevalence of VALGΦ8 in natural *V. alginolyticus* isolates (Chibani, Hertel et al. 2020) suggests that this phage may provide a selective advantage outside the laboratory in its natural environment, i.e., the pipefish. Conversely, however, bacterial genomes are graveyards of defective prophages (Bobay, Touchon et al. 2014), including filamentous phages (Davis, Moyer et al. 2000), suggesting that decay, rather than a peaceful coexistence, may be a common outcome for phages integrated into bacterial genomes. Ultimately, their level of virulence will dictate the fate of filamentous phages: Whereas lower virulence variants may enter into stable co-existence, higher virulence variants will be more prone to resistance-driven extinction and mutational decay if they do not provide a selective advantage.

## MATERIAL AND METHODS

Experiments were conducted using the *Vibrio alginolyticus* strain K01M1 (Chibani, Roth et al. 2020). K01M1 contains one integrated filamentous *Vibrio* phage VALGΦ6 (later called: resident K01M1Φ-phage throughout the manuscript), which replicates at a very low frequency (Chibani, Hertel et al. 2020). We compared VALGΦ6 and VALGΦ8 during previous work and found that both phages share relatively little sequence similarity (50.72%), except for proteins involved in DNA replication (Chibani, Hertel et al. 2020) and that VALGΦ6 does not confer SIE to VALGΦ8 (Wendling, Piecyk et al. 2017). Compared to other, closely related *V. alginolyticus* strains, K01M1 is highly susceptible to infections by filamentous phages, including VALGΦ8 (Wendling, Piecyk et al. 2017). For the selection experiment we used two different isogenic versions of the filamentous *Vibrio* phage VALGΦ8: VALGΦ8_K04M1_ (lower virulence) and VALGΦ8_K04M5_ (higher virulence; Table 1) which have been isolated from two different hosts (*V. alginolyticus* K04M1 and *V. alginolyticus* K04M5). The main difference between these two VALGΦ8 variants lies within the intergenic region upstream of gene K04M5_41300, whose function and impact we cannot explain (Figure S1). While both phages have been shown to significantly reduce the growth of K01M1 (Wendling, Piecyk et al. 2017, Wendling, Goehlich et al. 2018), infections with the higher virulence VALGΦ8_K04M5_ impose a significantly stronger reduction in bacterial growth than infections with the low virulence phage VALGΦ8_K04M1_. All experiments were carried out in liquid medium (Medium101: 0.5% (w/v) peptone, 0.3% (w/v) meat extract, 3.0% (w/v) NaCl in MilliQ water) at 25° C in 30-ml microcosms containing 6 ml of medium with constant shaking at 180 rpm.

### (a) Selection experiment

Six replicate populations were founded for each of three treatments from independent clones of K01M1. Treatments comprised (a) the higher virulence VALGΦ8_K04M1_, (b) the lower virulence VALGΦ8_K04M5,_ or (c) no phage as control. Each population was established from 60 *µ*l of an independent overnight culture (5×10^8^ CFU/ml). At the beginning of the experiment, we inoculated phage-containing treatments with 300 *µ*l of a 5×10^10^ PFU/ml stock solution. Populations were propagated by transferring 0.1% to fresh medium every 24 hours for a total of 30 transfers. On transfer T0, T1, T2 followed by every other transfer, phage and bacterial densities were determined, as described below and whole population samples were frozen at −80° C at a final concentration of 33% glycerol. In addition, on transfer T0, T1, T2, T6, followed by every sixth transfer 24 single colonies were isolated at random from each population and stored at −80° C. These colonies were later used during follow-up assays, as described below. Two populations from the control treatment tested positive for phage infection, indicating contamination, were excluded from later assays.

### (b) Bacterial and phage densities

*Bacterial densities*: bacterial densities were determined by plating out 100 *µ*l of a dilution series ranging from 10^−5^ to 10^−7^ on *Vibrio* selective Thiosulfate Citrate Bile Sucrose Agar (TCBS) plates (Sigma Aldrich). Plates were incubated over night at 25° C and the total amount of colonies was counted the following day.

*Phage densities*: quantification of filamentous phages by standard spot assays is often not possible (Rakonjac, Bennett et al. 2011). Instead of typical lytic plaques we mostly observed opaque zonas of reduced growth. Thus, we used spectrometry to quantify phage prevalence (http://www.abdesignlabs.com/technical-resources/bacteriophage-spectrophotometry), which uses the constant relationship between the length of viral DNA and the amount of the major coat protein VIII of filamentous phages, which, together, are the main contributors of the absorption spectrum in the UV range. The amount of phage particles per ml can be calculated according to the following formula:

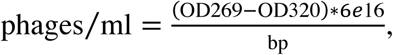

where OD269 and OD320 stand for optical density at 269 and 320 nm and bp stands for number of base pairs per phage.

The method is based on small-scale precipitation of phages by single PEG-precipitation. After centrifuging 1500 *µ*l of the phage containing overnight culture at 13,000 ×g for 2 min, 1200 *µ*l of the supernatant was mixed with 300 *µ*l PEG/NaCl 5× and incubated on ice for 30 min. Afterwards phage particles were pelleted by two rounds of centrifugation at 13,000 ×g for 2 min, resuspended in 120 *µ*l TBS 1× and incubated on ice. After one hour the suspension was cleaned by centrifugation at 13,000 ×g for 1 min and absorbance was measured at 269 and 320 nm.

Quantification of filamentous phages using spectrometry is likely to be erroneous if viral load is low. Therefore, we additionally quantified phage prevalence/ phage extinction in each of the populations on every second transfer day by standard spot assays with a serial dilution (up to 10^−6^) on the ancestral host (for details see (Wendling, Piecyk et al. 2017)) and measured until which dilution the typical opaque zones of reduced bacterial growth were visible.

### (c) Measuring phage-defence

We quantified the bacteria could not get infected by the respective ancestral phage by determining the reduction in bacterial growth rate (RBG) imposed by the phage, adapted from (Poullain, Gandon et al. 2008) with some modifications according to (Goehlich, Roth et al. 2019). Twenty-four random colonies from each population from transfer T0, T1, T2, T6, T12, T18, T24, and T30 were introduced into 96-well microtiter plates containing Medium101 at a concentration of 5×10^6^ cells/ml and inoculated with ∼2.5×10^6^ PFU/ml of the respective ancestral phage used for the selection experiment or without phage (control). Absorbance at 600 nm was measured using an automated plate reader (TECAN infinite M200) at T0 and again after 20 hours of static incubation at 25°C. The reduction in bacterial absorbance ‘RBG’ was calculated according to the following formula:

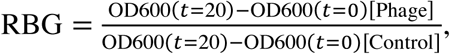

where OD stands for optical density at 600nm.

### (d) Frequency of prophage carriage

On transfer T0, T1, T2, T6 followed by every sixth transfer we measured the frequency of phage carriage of 24 random clones per population using standard PCR. We designed primers (VALGΦ8_Forward TGGAAGTGCCAAGGTTTGGT, VALGΦ8_Revers GAAGACCAGGTGGCGGTAAA) that specifically target the *Vibrio* phage VALGΦ8, but not the ancestral VALGΦ6, using the NCBI Primer-BLAST webpage (httdol://www.ncbi.nlm.nih.gov/tools/primer-blast/). Note, these primers only detect the presence of VALGΦ8, but not whether it exists episomally or as a prophage integrated into the chromosome. Glycerol stocks were inoculated overnight (25°C, 180 rpm) in Medium101 and subsequently diluted (1:10) in HPLC purified H_2_O and frozen at −80° C. One *µ*l of this suspension was used as DNA template in the PCR assay. Reaction comprised 1 *µ*l Dream Tag Buffer, 0.1 *µ*l Dream Tag DNA polymerase (Thermo Scientific, USA), 4.9 *µ*l H_2_O, 1 *µ*l dNTPs [5 mM] and 1 *µ*l of each primer [50 *µ*M]. The amplification program used consisted of: (i) 3 min at 95° C, (ii) 35 cycles of 45 sec at 95° C, 30 sec at 63° C, 45 sec at 72° C, (iii) 7 min at 72° C. Afterwards, 5 *µ*l of each reaction was mixed with 2 *µ*l loading dye (10×) and loaded onto a 1.2% agarose gel dissolved in 1×TAE gel buffer. GeneRuler Express DNA-ladder was used as size marker. Agarose gels were run 15 min at 70 V in 0.5× TAE running buffer and subsequently stained with ethidium bromide for 10 min. DNA was visualized using UV light and documentation took place using Intas Gel iX20 Imager. Phage presence was recorded positive if a PCR product of 1209 bp was visible.

For all subsequent assays, we randomly picked one immune clone with a positive PCR product (later called: super infection exclusion SIE hosts) and one resistant clone with a negative PCR product (later called: surface receptor mutant SRM host) from each phage-evolved population as well as two randomly selected non-resistant clones from the control populations.

### (e) Competition experiments

To determine differences in fitness between both resistance forms, we measured the competitive fitness of three randomly selected PCR positive relative to three randomly selected PCR negative clones from each treatment. Each competition culture was done in triplicates as described in (Harrison, Guymer et al. 2015). In brief, overnight cultures of both competing strains (of which one was labelled with a GFP-marker) were mixed 1:1 and 60 *µ*l of this mixture was inoculated to 6 ml Medium 101 to initiate each competitive culture. After 24 hours, fitness was estimated by means of flow cytometry (FACS-Caliburm Becton & Dickinson, Heidelberg, GER), where absolute fluorescent cells and non-fluorescent cells were calculated. Competitive fitness was estimated as the ratio in Malthusian parameters (Lenski, Rose et al. 1991):

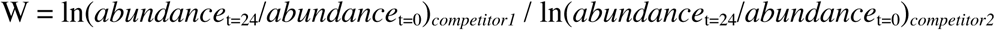

### (f) Bacterial growth rate and phage production

To determine fitness parameters that could explain observed differences in competitive fitness we additionally quantified bacterial growth rate (*µ*) by means of 24-hour growth curves and phage production using PEG precipitation (as described in (c)) of the same clones used for the competition assays (i.e., one SIE and one SRM host from each phage-treated population and two random phage-susceptible clones from the control populations plus ancestors).

### (g) Motility

Motility was visualized on mid-exponential growth cultures using a light microscope and swimming was captured for 50s.

### (h) Whole genome sequencing

We used a combination of long- and short read sequencing to obtain complete genomes of the same clones from the assays above, i.e., one SIE and one SRM host from each phage-treated population and one random phage-susceptible clone from each control population, which corresponds to six independently evolved clones per treatment and resistance form. Clones were taken from timepoint 2, because phage-carriers disappeared quickly from the populations and we were thus not able to pick one mutant and one SIE host from the same timepoint and population later than timepoint two. High molecular weight DNA was extracted from cell pellets of overnight cultures following the protocol for gram negative bacteria from the DNeasy Blood & Tissue Kit (Qiagen, Hilden, Germany). Long-read sequencing was performed at the Norwegian Sequencing Centre according to the following protocol: the library was prepared using Pacific Bioscience protocol for SMRTbell™ Libraries using PacBio^®^ Barcoded Adapters for Multiplex SMRT^®^ Sequencing. To do so, DNA was fragmented into 10kb fragments using g-tubes (Covaris). Samples were pooled during library preparation aiming for equimolar pooling and library size was selected using Ampure beads. The library was sequenced on a PacBio Sequel instrument using Sequel Polymerase v3.9, SMRT cells v3 LR and Sequencing chemistry v3.0. Loading was performed by diffusion. Two SMRT cells were sequenced (movie time: 600min, pre-extension time: 240 min). Reads were demultiplexed using Barcoding pipeline on SMRT Link (v6.0.0.47841, SMRT Link Analysis Services and GUI v6.0.0.47836) with 40 as a minimum barcode score.

For short read sequencing concentration and purity of the isolated DNA was first checked with a Nanodrop ND-1000 (PeqLab Erlangen, Germany) and exact concentration was determined using the Qubit® dsDNA HS Assay Kit as recommended by the manufacturer (Life Technologies GmbH, Darmstadt, Germany). Illumina shotgun libraries were prepared using the Nextera XT DNA Sample Preparation Kit. To assess quality and size of the libraries, samples were run on an Agilent Bioanalyzer 2100 using an Agilent High Sensitivity DNA Kit as recommended by the manufacturer (Agilent Technologies, Waldbronn, Germany). Concentration of the libraries were determined using the Qubit® dsDNA HS Assay Kit as recommended by the manufacturer (Life Technologies GmbH, Darmstadt, Germany). Sequencing was performed on a MiSeq system with the reagent kit v3 with 600 cycles (Illumina, San Diego, CA, USA) as recommended by the manufacturer and resulted in a minimum average coverage of 88× per strain (coverage range was from 88× to 157×). The reads were quality controlled using the program FastQC Version 0.11.5. All illumina reads that passed the FastQC filtering were used for hybrid assemblies as well as for single nucleotide variation analysis.

Genome assemblies were performed in two different ways: (i) long-read data was generated for all replicates where the presence of the infecting phage was confirmed by PCR. The Assemblies were performed as hybrid assemblies using short-read and long read data in a short-read first approach. In brief: An initial assembly was performed with short-read only using spades (v3.13.0) as provided within Unicycler (Bankevich, Nurk et al. 2012). The resulting contigs were co-assembled with long-read data using miniasm (v0.2-r168) (Li 2016) and curated using the racon software (Vaser, Sovic et al. 2017). This step resulted in complete closed replicons. All long reads were mapped and integrated into the contigs. All replicons were polished using Pilon (v1.22) to clear any small-scale assembly errors (Walker, Abeel et al. 2014). Finally, all replicons were rearranged according to the origin of replication. (ii) the assembly for the ancestral K01M1 strain, as has been described in (Wendling, Piecyk et al. 2017) was performed following the Hierarchical Genome Assembly Process (HGAP3) protocol, developed for Pacific Biosciences Single Molecule Real-Time (SMRT) sequencing data (Chin, Alexander et al. 2013). HGAP is available for use within PacBio’s Secondary Analysis Software SMRTPortal. Methodically, the longest subreads of a single SMRT Cell (usually 25x genome coverage, e.g., 25 × 5 Mbp = 125 Mbp) are being chosen to be error-corrected with “shorter” long reads in a process named preassembly. Hereby, a length cut-off is computed automatically separating the “longer” reads (for genome assembly) and the “shorter” reads (for error-correction). The level of error-correction is being estimated with a per-read accuracy of 99%. Finally, error-corrected long read data is being assembled with Celera Assembler (v7.0) (Langmead and Salzberg 2012).

### (i) SNV analysis and reconstruction of infecting phages

All short-read sequences were mapped on a high quality closed reference genome of *Vibrio alginolyticus* Strain K01M1 (14) using Bowtie2 (Langmead and Salzberg 2012). Single nucleotide variation (SNV) analysis was done using the Breseq pipeline as described in (Deatherage and Barrick 2014). To ensure that observed sequence variations in resistant phenotypes are real mutations and no read-mapping errors we filtered out all variants that occurred in the control treatment.

We calculated whole genome alignments of evolved clones using the MAUVE aligner (Darling, Mau et al. 2010). Presence of infecting phage genomes were confirmed by assembling NGS-reads that did not map on the K01M1 genome in a bowtie2 mapping using Spades (Bankevich, Nurk et al. 2012). The resulting contigs were annotated based on the review (Mai-Prochnow, Hui et al. 2015) on filamentou s phages. The genomes of the evolved phages were compared to the infecting phage genomes *Vibrio* phage VALGΦ8 as well as to the genome of the resident prophage *Vibrio* phage VALGΦ6 from the challenged strain K01M1 using BLAST and Easyfig 2.1 (Sullivan, Petty et al. 2011). We found no evidence for chromosomal integration of VALG Φ8. Instead, we found multiple lines of evidence for an episomal existence of VALGΦ8 in all PCR+ clones:

i. we mapped the illumina reads of the ancestral K01M1 genome and generated a fastq file from all reads that did not map. Next we assembled these non-mapping reads using Spades (Bankevich, Nurk et al. 2012) and analysed the resulting assembly. We found individual contigs that consisted of the complete genome sequence of VALGΦ8 (Figure S3) with overlapping ends. From this we concluded that a circular VALGΦ8 genome representing an episomally existing phage can be reconstructed from illumina reads.
ii. we found no hybrid reads that contain both bacterial and VALGΦ8 DNA, and concluded that this phage did not integrate into the chromosome
iii. we found no new junction events in our breseq analysis containing both phage and bacterial sequences which supports our conclusion that VALGΦ8 did not integrate into the bacterial chromosome
iv. whole genome alignments from hybrid-assemblies using PacBio and illumina reads using the unicycler software with the ancestral K01M1 genome revealed no structural variants. This again supports the conclusion that there are no integrated VALGΦ8 prophages present in the evolved probes.
v. an in-detail investigation of known Inoviridae integration loci (e.g., t-RNAs, pilus genes) using the assemblies of the PCR positive clones revealed that none of these loci contained VALGΦ8 which supports again our conclusion that VALGΦ8 did not integrate into the chromosome.
vi. from the PacBio long-read data we found reads that cover multimers of the complete ancestral VALGΦ8 and concluded from there that the episomally existing VALGΦ8 replicates via the rolling circle mechanism

### (j) Statistical analysis

All statistics were performed in the R 4.0.4 statistical environment. For all analysis aimed to compare the two different phage treatments to one another, control populations (i.e., those that evolved without phages) were excluded. When comparing temporal dynamics between phage-treatments, we excluded the starting time-point T0, because these measurements were taken before phages were added to the populations.

#### Bacteria and phage dynamics

Bacterial and phage densities were analysed over time using a generalized least squares model to control for autocorrelation of residuals over time using the gls function (package nlme) with phage treatment, transfer as categorical variable as well as their interaction as fixed effect.

We considered phages to be prevalent in the population if opaque zones of reduced growth were visible during standard spot assays. Phage prevalence was subsequently quantified by a serial dilution, which were assigned with invers values (i.e., if reduced growth zones were visible up to dilution of 10^−6^ we assigned to it a value of 7, whereas if they were only visible on an undiluted spot, we assigned to it a value of 1, if no zone of reduced growth was visible it was scored as 0). Phage extinction events across phage-treatments were analysed using a log-rank test.

#### Measuring phage defence and prevalence

We observed a bimodal histogram on all RBG values with a local minimum at RBG = 0.82 (Figure S2). Thus, we considered an infection as positive if RBG < 0.82. The proportion of clones per population that could not get infected by the ancestral phage as well as the proportion of clones that tested positive for PCR (targeting the VALGΦ8) were analysed using a generalized linear model with a binomial error distribution using the glm function (package lme4) with phage treatment, transfer and their interaction as fixed effect.

#### Fitness effects

We determined differences in relative fitness between SRM and SIE hosts using a linear model with resistance mechanisms and GFP-label and the interaction thereof as fixed effects. Maximum growth rates (*µ*) were estimated for each strain by fitting generalized logistic models to individual growth curves using the open-source software package Curveball (https://github.com/yoavram/curveball) (Ram, Dellus-Gur et al. 2019) in python. To determine differences in the amount of free phages and in growth rates produced between ancestral strains and evolved strains and between both resistance forms, we used Welch’s pairwise *t*-tests with sequential Bonferroni correction. We further performed a Pearson’s correlation analysis to determine whether phage production impacted bacterial growth rates.

### (k) Mathematical model

We modelled the dynamics of the non-resistant evolved clones (with density *B*), resistant SIE hosts (*I*), resistant SRM hosts (*R*), and SIE hosts that have also acquired the MSHA mutation (*IR*), as well as the phage population (*V*) in batch cultures by the following system of differential equations:

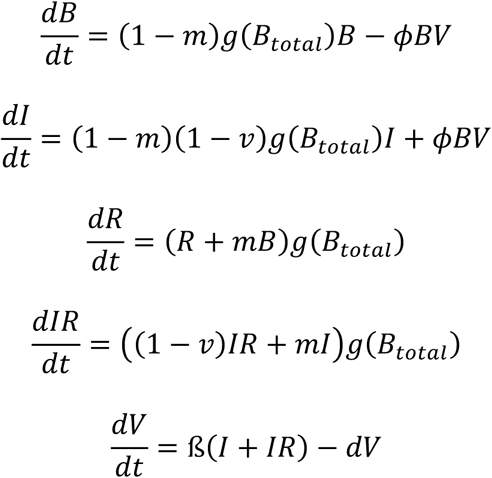

Bacterial growth was modelled by generalized logistic growth of the form 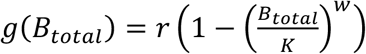. Here *r* is the maximum growth rate (mgr) of the non-resistant evolved bacteria, *K* is the carrying capacity of the batch culture and *B*_*total*_ = *B* + *I* + *R* + *IR* is the total density of all bacterial types. The curvature parameter *w* determines whether maximum growth is attained at an early point in the growth phase (*w* < 1) or at a late point (w > 1). We assume that SIE hosts (I and IR) suffer a growth rate reduction relative to the non-resistant evolved bacteria due to virulence caused by intra-cellular production of virus particles, here represented by the virulence factor v <= 1. A completely avirulent phage would have v=0, and maximum virulence v=1 corresponds to growth arrest of the bacterial cell.

Phages (*V*) infect non-resistant evolved bacteria (*B)* following a mass action law with adsorption rate (phi), reflecting that increasing densities of either bacteria or phages lead to higher encounter rates and thus more infections. Infection of a bacterial cell transforms cells into a resistant phage carrier, SIE host (*I)*, which actively produces new viral particles (*V*) with phage production rate (*ß*). Additionally, both non-resistant evolved bacteria (*B)* and SIE hosts (*I)* can acquire complete resistance (*R & IR*) through mutations within the MSHA type IV pilus operon. We assume that SRM hosts have the same growth rate as the non-resistant evolved bacteria.

All bacterial types grow until the carrying capacity (*K*) is reached, but bacteria-phage interactions continue to occur as long as there are sensitive bacteria and phages left. After a certain time t*max* a portion (here 1/1000th) of the entire community is transferred to fresh medium and the process restarts.

## Supporting information

Supplementary video material ancestor

Supplementary video material SRM host

## Acknowledgements

We thank Pratheeba Pandiaraj, Katja Trübenbach, Veronique Merten, Silke-Mareike Merten and Kim-Sara Wagner for their support in the laboratory.

## Data availability

All experimental data have been deposited on dryad (a link will be provided upon acceptance of the manuscript). Genomic data is available at NCBI (accession number will be provided upon acceptance of the manuscript), and in the supplemental data file Table S1 and S2.

