## Supplementary material for "Higher phage virulence accelerates the evolution of host resistance": Table S1-S3, Figure S1- S4

### Tables

**Table S1** SNV Table of all clones evolved in the presence of either the higher virulent (ΦK04M5) or the lower virulent (ΦK04M1) phage VALGΦ8

| Gene | Strain/Position | Mutation | Annotation | Phage |
|---|---|---|---|---|
| K01M1_00270<br>/ K01M1_00280 | 27910<br>28000<br>28036<br>28093 (*) | G→A<br>A→T<br>A→T<br>+T | tRNA-Val(23S rRNA intergenic | ΦK04M5<br>ΦK04M1 |
| K01M102170 | **VALG24:**<br>230316 | G→T | hypothetical protein | ΦK04M5 |
| K01M1_18150<br>/ K01M1_18160<br>/ K01M1_18170<br>/ K01M1_18180<br>/ K01M1_18220<br>/ K01M1_18230 | **Different replicates (*)** | +A<br>(A)6→5<br>+G<br>+A<br>Δ1 bp | hypothetical protein | ΦK04M5<br>ΦK04M1 |
| mshL | **VALG25**:<br>3024691 | G→A | putative secretin T2SS | ΦK04M5 |
| K01M1_28150 | **VALG26:**<br>3012432<br>**VALG33**:<br>3014427<br>**VALG32**:<br>3014807<br><br><br>**VALG28**:<br>3015059 | (Δ9)<br><br>(+T) → frameshift<br><br>(duplication of<br>CTCGTTATTACAGATATA)<br><br><br>TAGAGA 4 → 5 | putative secretin, similar to protein D of T2SS, protein G of T3SS outer membrane component | ΦK04M5<br>ΦK04M1<br>ΦK04M1<br><br><br>ΦK04M5 |
| mshE | **VALG27:**<br>3021814<br>**VALG24:**<br>3022029 | Δ11 bp → frameshift<br><br>+C → frameshift | P-loop NTPase, component of T2SS and T4SS, interacts with mshG | ΦK04M5 |
| mshG | **VALG31:**<br>3020189 | (GGTGAT) duplication | Inner membranre component of T2SS and T4SS, interacts with mshE | ΦK04M1 |
| cadB/cadC intergenic | **VALG29**:<br>3243856 | (TAGAGA)4 → 5 | putative cadaverine/lysine antiporter/Transcriptional activator CadC | ΦK04M1 |

(*) at least one of the mutations listed for this genome region occurred in every infected clone

**Table S2** Number of reads that map uniquely to VALGΦ8 from each sequenced clone. Read counts on PCR-negative clones are derived from sequences of the resident VALGΦ6 located in the chromosome of strain K01M1, see Figure S4.

| Sample | total read pairs | matches on VALGΦ8 | PCR-positive |
|---|---|---|---|
| VALG11 | 984193 | 94301 | + |
| VALG12 | 1080448 | 685 | + |
| VALG13 | 1285444 | 9498 | + |
| VALG14 | 1669371 | 306 | + |
| VALG15 | 1349048 | 283 | - |
| VALG16 | 1320275 | 20125 | + |
| VALG17 | 1456946 | 3300 | + |
| VALG18 | 1467665 | 6308 | + |
| VALG19 | 1574017 | 38691 | + |
| VALG20 | 1269481 | 1359 | + |
| VALG21 | 1142477 | 60215 | + |
| VALG22 | 1342309 | 1345 | + |
| VALG23 | 1211521 | 177 | - |
| VALG24 | 1592216 | 340 | - |
| VALG25 | 1283594 | 206 | - |
| VALG26 | 1564912 | 18315 | + |
| VALG27 | 1332438 | 232 | - |
| VALG28 | 1373230 | 232 | - |
| VALG29 | 1449256 | 269 | - |
| VALG30 | 1608391 | 277 | - |
| VALG31 | 1378663 | 213 | - |
| VALG32 | 1161115 | 219 | - |
| VALG33 | 1122114 | 203 | - |
| VALG34 | 1771100 | 332 | - |
| VALG47 | 1693657 | 372 | - |
| VALG48 | 1965837 | 493 | - |
| VALG49 | 1237655 | 264 | - |
| VALG50 | 1394821 | 329 | - |
| VALG51 | 1460129 | 369 | - |
| VALG52 | 1731878 | 434 | - |

**Table S3:** ANOVA table competition assay between SIE hosts and MSHA-mutants. Treatment comprises lower or higher virulence phage. Assays have been performed twice, where either the SIE hosts or MSHA-mutants were carrying the GFP marker.

| | Df | Sum Sq | Mean Sq | F value | Pr(>F) |
|---|---|---|---|---|---|
| Treatment | 1 | 0.47716 | 0.47716 | 18.6292 | 0.002559 |
| GFP Label | 1 | 0.10144 | 0.10144 | 3.9602 | 0.081770 |
| Treatment:Label | 1 | 0.00035 | 0.00035 | 0.0137 | 0.909871 |
| Residuals | 8 | 0.20491 | 0.02561 | | |

### Figures

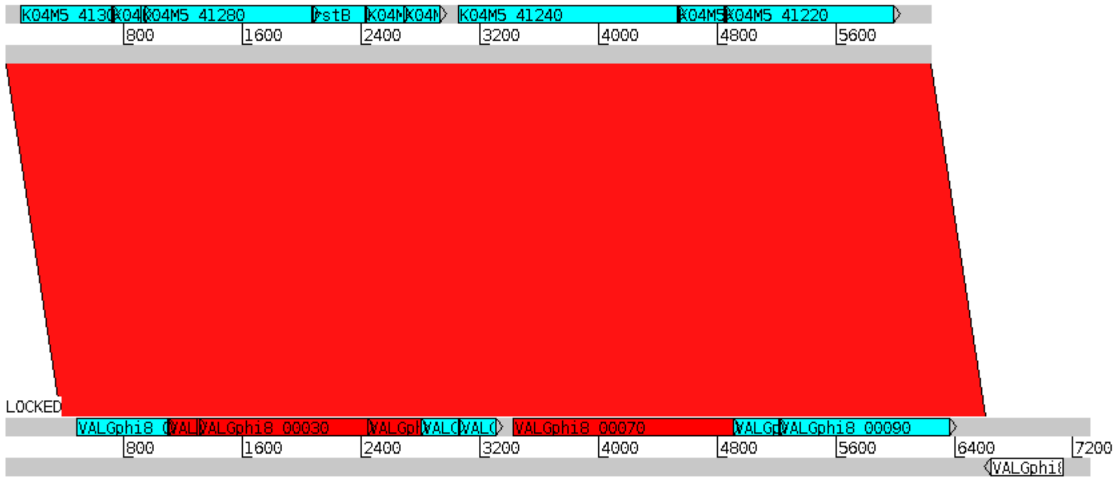


**Figure S1** Alignment between higher (top) and lower virulent (bottom) version of VALGΦ8 (A). The genomes of the two VALGΦ8 phages are identical within all coding regions. The only sequence difference between the genomes has been identified within the intergenic region upstream of gene K04M5_41300.

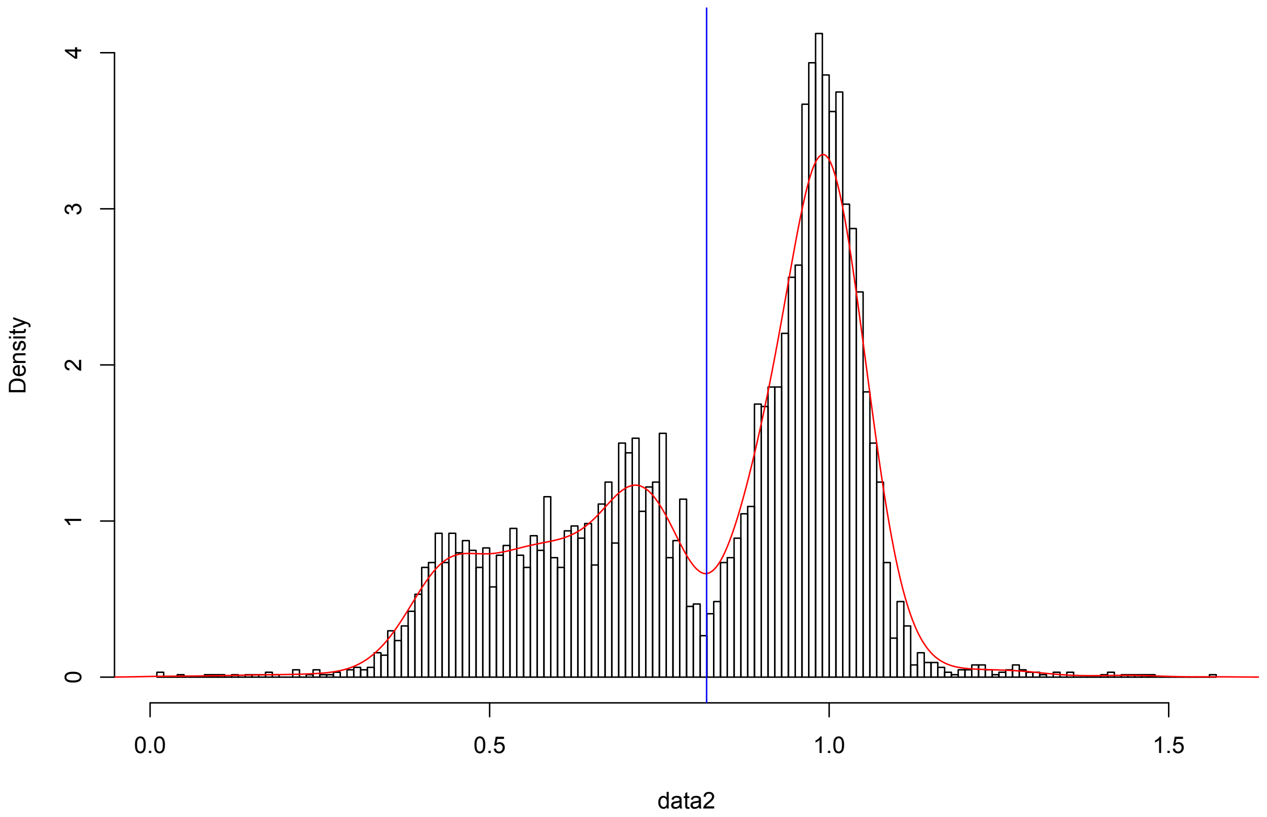


**Figure S2 Frequency of RBG-values**. From the bi-modal distribution we calculated the local minimum at 0.82 and concluded that all RBG-values < 0.82 indicate a significant reduction in bacterial growth imposed by the phage.

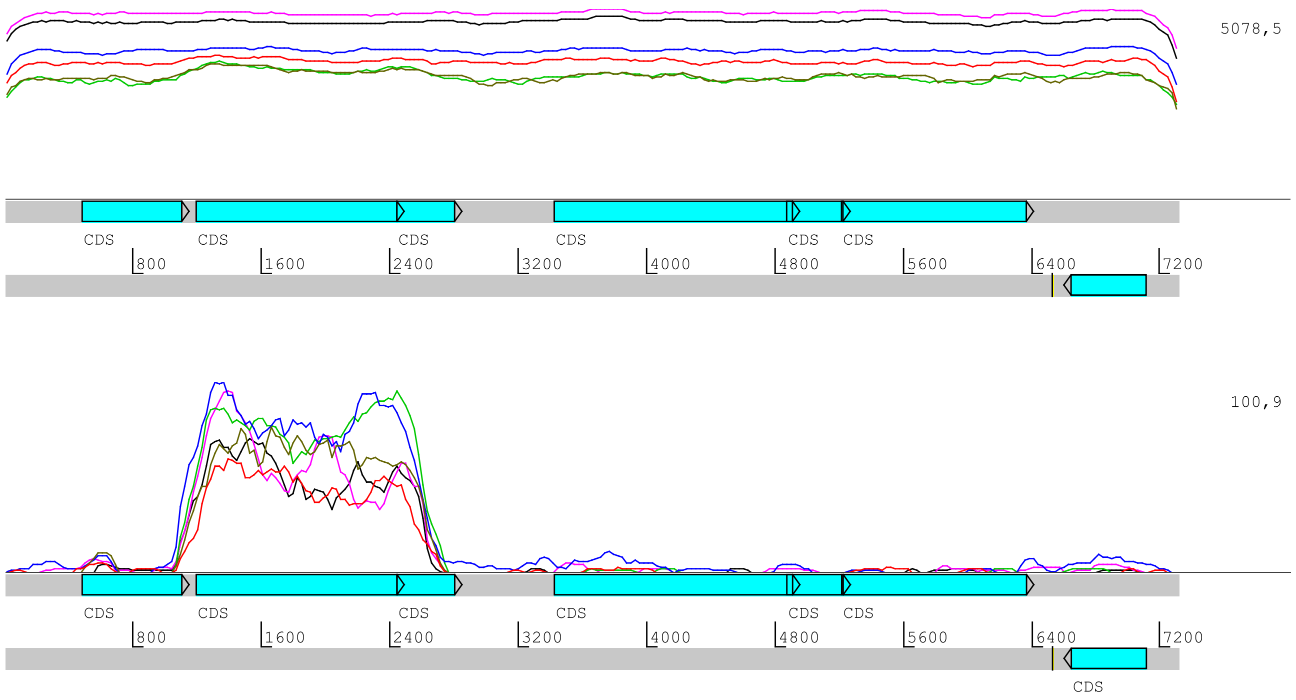


**Figure S3**: Reads mapping uniquely to VALGΦ8. Top: PCR-positive clones, reads span the entire region of the infecting VALGΦ8. Bottom: PCR-negative Clones, here VALGΦ6 reads map to a shared region between the ancestral VALGΦ6 and the infecting VALGΦ8.

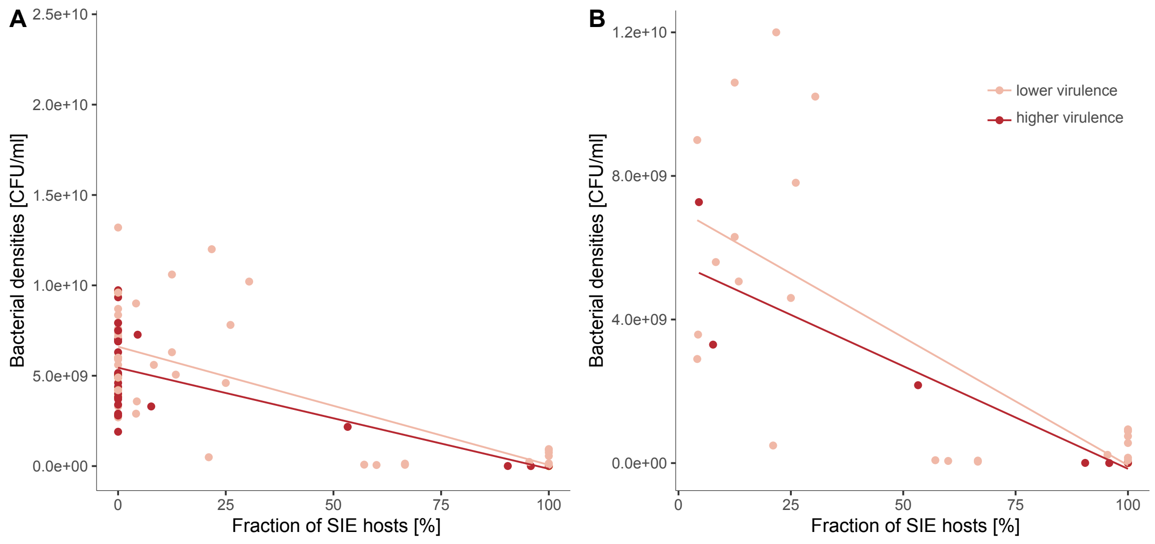


**Figure S4**: Correlation between the fraction of SIE hosts per resistant clones and total bacterial population size measured as CFU/mL (left with zero inflation, right: without zero inflation. Higher virulence phage VALGΦ8$_{K04M5}$ (dark red), lower virulence phage VALGΦ8$_{K04M1}$ (light red).
